# Differentiation and migration of hematopoietic stem and progenitor cells cross multiple tissues

**DOI:** 10.1101/2023.09.15.557856

**Authors:** Shiya Yu, Hui Li, Xuefei Wang, Guanming Chen, Hongwen Huang, Ni Hong, You-Qiang Song, Xuegong Zhang, Wenfei Jin

**Affiliations:** School of Life Sciences, Southern University of Science and Technology, Shenzhen, China; School of Biomedical Sciences, The University of Hong Kong, Hong Kong, China; MOE Key Lab of Bioinformatics and Bioinformatics Division of TNLIST, Department of Automation, Tsinghua University, Beijing, China; CAS Key Laboratory of Computational Biology, Shanghai Institute of Nutrition and Health, University of Chinese Academy of Sciences, Chinese Academy of Sciences, Shanghai, 200031, China

**Author notes:** These authors contributed equally (Shiya Yu, Hui Li). Corresponding author (W.J.).

**Keywords:** Single cell sequencing, Cell migration, Differentiation cross tissues, Hematopoiesis, Hematopoietic stem cells and progenitor cells (HSPCs)

## Abstract

Hematopoiesis requires the coordinated differentiation of hematopoietic stem cells and progenitor cells (HSPCs) in multiple tissues. Although differentiation of HSPCs in bone marrow (BM) has been well-studied, our knowledge about the migration and differentiation of HSPCs cross tissues is limited. Here, we collected and integrated single-cell RNA-seq data of human CD34+ cells, which represent HSPCs, from BM, peripheral blood (PB), thymus and mobilized PB (mPB), to investigate the hematopoiesis cross tissues. We constructed a cell atlas of HSPCs cross tissues and found most HSPC subsets in BM had counterparts in PB, indicating migration of HSPCs from BM to PB has a much broad spectrum. We found B progenitors highly expressed *CXCR4* for anchoring in BM, while cells with low expression of *CXCR4* facilitate their migration out of BM. Among the HSPC subsets from thymus, we only found the counterparts of the earliest thymic progenitors (ETPs) in BM and PB, potentially indicating that ETPs were the subsets that migrated from BM to PB and thymus. We found interaction signaling including *CD99*-*CD99*, *CXCL12*-*CXCR4* and *CCL19*-*CCR7* played important roles in ETP homing to thymus. Briefly, these data provided a single unified developmental spectrum of hematopoiesis cross different tissues, connected by cell migration.

## Introduction

Human blood is the most regenerative tissue, with approximately one trillion cells produced through hematopoiesis daily. Hematopoiesis generates immune cells through the differentiation of a small number of self-renewing multipotent hematopoietic stem cells (HSCs) into various progenitor cells^1–4^. Early studies showed hematopoiesis occurs through a stepwise process from pluripotent progenitors, to oligopotent progenitors, to unipotent progenitors, terminating in mature blood cells^5,6^. In recent decade, massively parallel single-cell RNA-seq (scRNA-seq) enabled routine analyses of thousands of or even tens of thousands single cells for inferring the developmental lineages. Single cell analysis of hematopoietic stem and progenitor cells (HSPCs) from bone marrow (BM) demonstrated hematopoiesis is a continuously hierarchical process^4,7–10^. The dynamics of gene expression, lineage and developing commitment during hematopoiesis has to be precisely regulated^10–12^. Dysregulation or malfunction of hematopoiesis may lead to various diseases such as myelofibrosis (MF)^13^, leukemias^14^ and sickle cell disease^15^.

There are many studies on hematopoiesis in BM, which showed that hematopoiesis is a continuously hierarchical process^4,8,10,14,15^. Popescu *et al*.^16^ analyzed hematopoiesis in human fetal liver and found a shift in haemopoietic composition of fetal liver during gestation. Zheng *et al.*^7^ analyzed the HSPCs in human cord blood to identify the molecular transitions during early hematopoiesis. Mende *et al.*^17^ studied unique molecular and functional features of human extramedullary HSPCs. Psaila *et al.*^13^ analyzed mobilized PB (mPB) CD34+ cells in healthy individuals and revealed megakaryocyte-biased hematopoiesis in MF patients. Le *et al.*^18^ analyzed CD34+ cells in thymus and revealed multilineage priming followed by gradual commitment to T cell lineage during initial stages of thymopoiesis. These studies provided biological insights into the process of hematopoiesis in one or several tissues. However, hematopoiesis is a complex process that requires multi-tissues coordination, and the mechanism of coordinated migration and differentiation of HSPCs among multi-tissue is still unclear.

To investigate the differentiation and migration of HSPCs, we obtained 90,598 single cell transcriptomes of CD34+ cells from four tissues, namely BM, PB, mPB and thymus. Our integrated analyses of CD34+ cells showed the hierarchical structure of hematopoiesis cross the tissues. We found almost all HSPC subsets in BM had counterparts in PB/mPB, indicating migration of HSPCs from BM to PB/mPB has a much broad spectrum. We further analyzed migration abilities of these HSPC subsets in different tissues, and inferred the signals underlying HSPCs remaining in BM or migration out of BM. Among the cell subsets from thymus, we only found the counterparts of the earliest thymic progenitors (ETPs) in PB and BM, indicating that ETPs were the subset that migrated from BM and thymus. We further analyzed the trajectory of ETPs and cell-cell interaction signaling for inferring the potential mechanism of ETPs homing to thymus.

## Results

### A single cell transcriptome atlas of human HSPCs in BM, PB, mPB and thymus

We designed a workflow to investigate the differentiation and migration of HSPCs cross multiple tissues (**Fig 1a**). In brief, by integrating scRNA-seq data of CD34+ cells from multiple tissues, we construct a comprehensive cell atlas of HSPCs and infer hematopoietic lineages. We further infer cell migration cross tissues by jointly considering the hematopoietic lineages and the tissues differences of each HSPC subset. We obtained total 90,598 single cell transcriptomes of CD34+ cells from BM^4,15^, PB^17^, mPB^13^ and thymus^18^ (**Supplementary Table 1**). We integrated these scRNA-seq data with Seurat ^19^ and projected these cells on Uniform Manifold Approximation and Projection (UMAP) (**Fig 1b**). The HSPCs almost formed a single connected entity on UMAP, which extended into 4 main branches and were grouped into 18 clusters (**Fig 1b**). We inferred the cell type of each cluster by checking their expression of HSC specific genes and hematopoietic lineage specific genes, including HSC (*MEG3*+, *AVP*+, *CRHBP*+, *Lin*-), multipotent progenitor (MPP) (*AVP*+, *CRHBP*+, *CSF3R*+), lymphoid-primed multi-potential progenitor (LMPP) (*CSF3R*+, *CRHBP*+), myeloid (My) progenitor (*MPO+*, *LYZ*+ and *IRF8*+), common lymphoid progenitor (CLP) (*CCR7*+, *NKG7*+), B progenitor (*VPRE*+, *EBF1*+), T progenitor (ETP, Thy #1, Thy #2, Thy #3) (*CD3D*+, *CD7*+, *RAG2*+, *CD1A*+), megakaryocytic (Mk) progenitor (*GATA2*+, *PF4*+, PPBP+) and erythroid (Ery) progenitor (*GATA2*+, *CD36*+) (**Fig 1c; Supplementary Fig 1a)**. Pseudotime analyses showed differentiation of HSCs following a continuous hierarchical process (**Fig 1d)**, in which HSCs differentiated into two MPP subsets, with MPP #1 differentiated into Ery lineage and Mk lineage, while MPP#2 differentiated into LMPP then immune related lineages including My lineage, B lineage and T lineage, consistent with previous studies^1–4,8–10^. Furthermore, the HSC score is decreasing along pseudotime trajectory (**Supplementary Fig 1b**).

**Fig 1.**
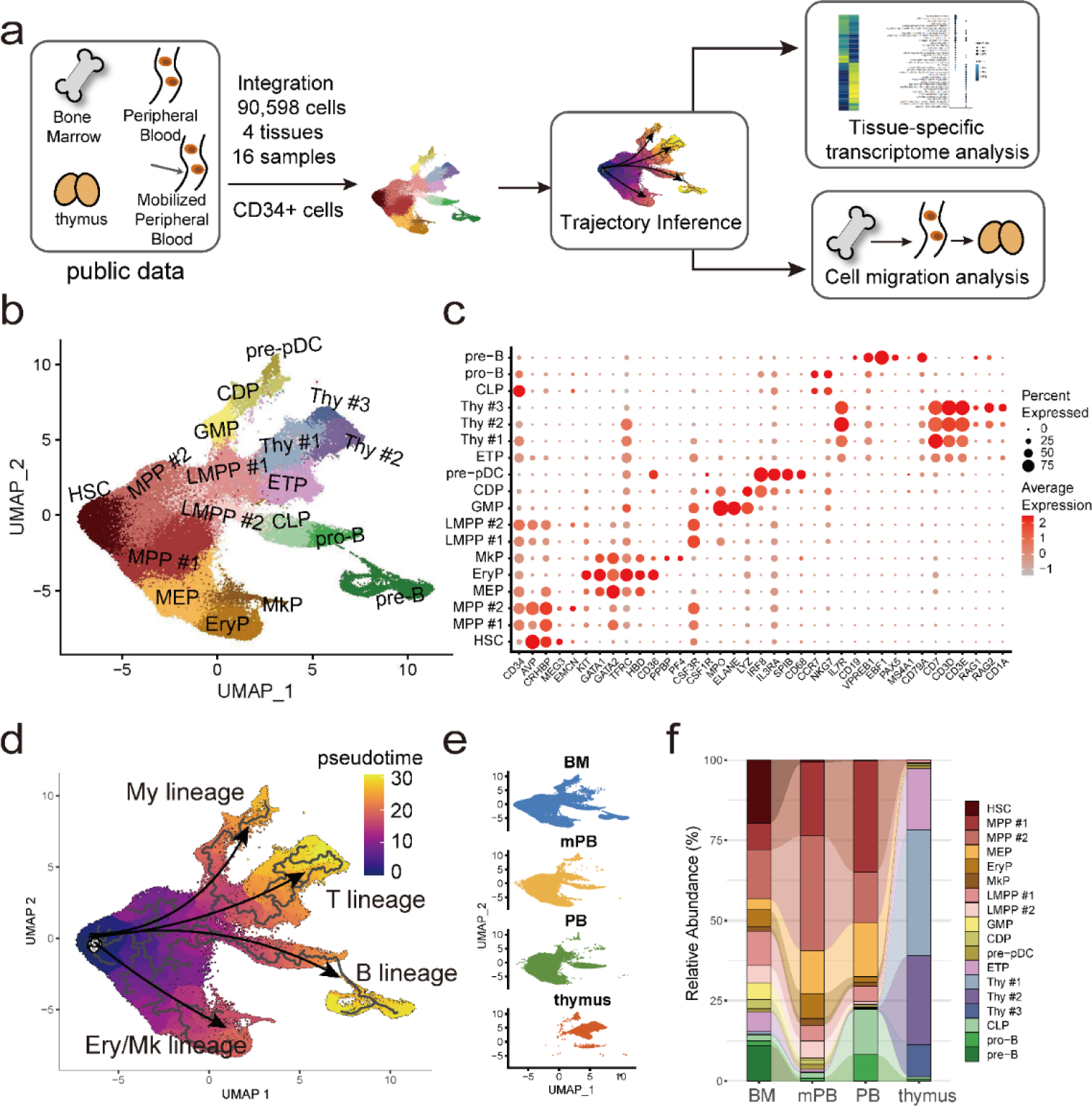
A single cell transcriptome atlas of human CD34+ cells in bone marrow (BM), peripheral blood (PB), mobilized PB and thymus. **a**. Scheme of this study. **b**. UMAP visualization of CD34+ cells in BM, PB, mPB and thymus, colored by inferred cell subset. **c**. Normalized expression level and expression percentage of the cell subset-specific genes in the 18 HSPC subsets. **d**. Hematopoietic lineages inferred by Monocle3. **e**. UMAP visualization of CD34+ cells in each tissue. **f**. Stacked bar plot of percentage of cell subset in each tissue.

We found that cells from BM, PB, and mPB had similar distribution on UMAP plot and are highly overlapped (**Fig 1e, f**). While most thymus CD34+ cells are in T lineage, with some fractions in My lineage and B lineage (**Fig 1e, f**), consistent with the report that early T progenitors in thymus have B and myeloid potential^18,20^. HSPCs from BM have the broadest distribution on UMAP and have the most complete HSPCs subsets including pre-B which is almost exclusively present in BM (**Fig 1e, f**). Thymus CD34+ cells only had a little overlap with BM and PB in the earliest stages of T progenitors, consistent with recent studies on thymus seeding progenitors (TSPs)^21^.

### Comparison of HSPCs between BM, PB and mPB

Although the hierarchical structure of HSPCs in BM, PB and mPB are similar (**Fig 2a**), the cell subsets from the three tissues showed some differences. In particular, the cells in HSC, GMP and pre-B are mainly from BM, while the cells in CLP and pro-B are mainly from PB (**Fig 2b**). The cells in LMPP#1, LMPP#2, CDP, pre-pDC, EryP and are mainly from BM and mPB, with a little contribution from PB (**Fig 2b**). We calculated the density of cells along the pseudotime in each tissue. The results showed that cells from BM dominated the early hematopoiesis and their cell density decreased gradually along the pseudotime (**Fig 2c**). Cells from PB and mPB were rare in the earliest hematopoiesis, with cell density increasing rapidly along the pseudotime and peaking at around one third of the pseudotime, then gradually decreasing (**Fig 2c**). Cell density along each hematopoietic lineage is similar to that based on overall pseudotime (**Fig 2c; Supplementary Fig 2a**). These results suggested that PB and mPB have similar cell subsets and have similar lineage features, both of which are quite different from that of BM.

**Fig 2.**
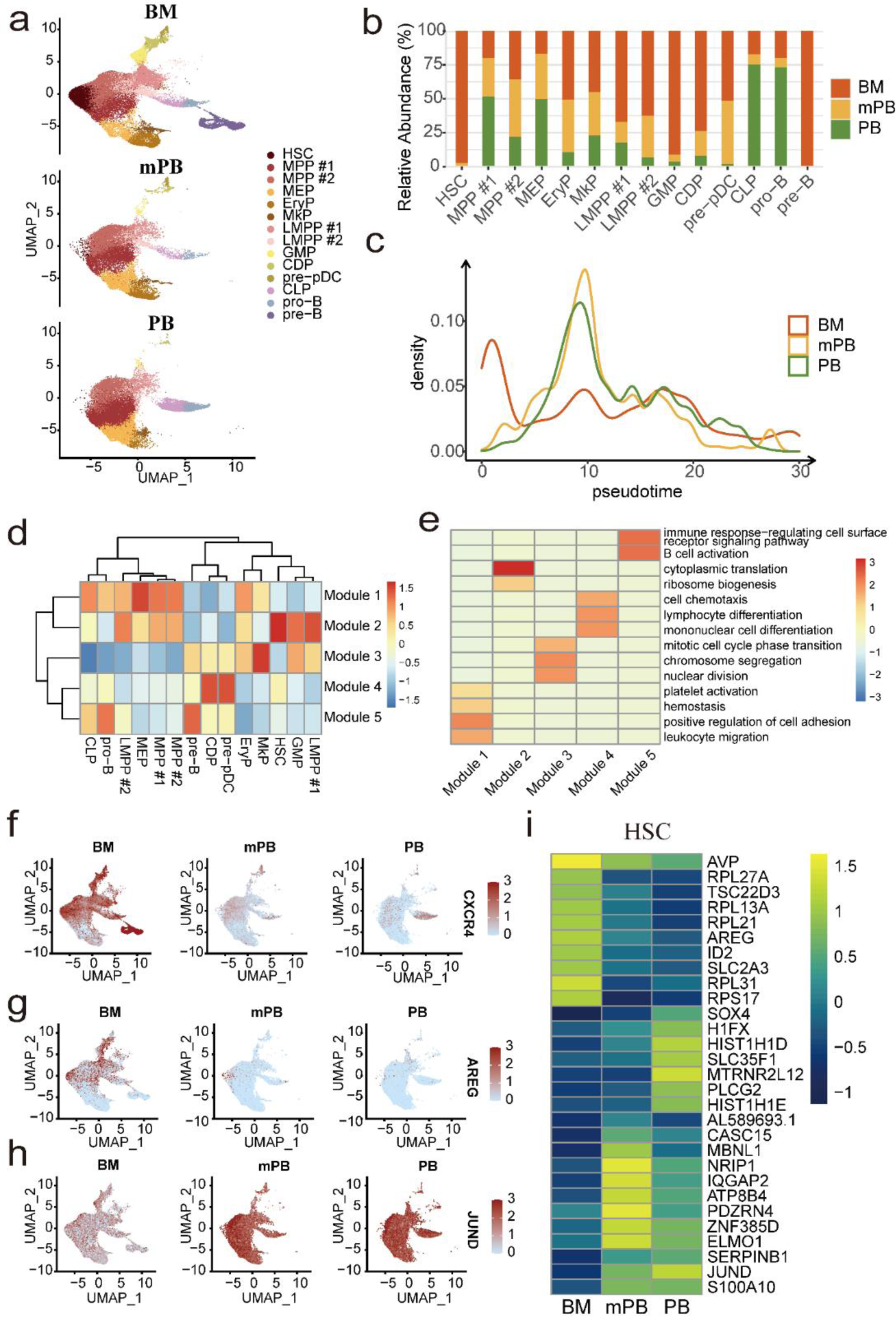
Comparison of hematopoietic lineage between BM, PB and mPB. **a**. UMAP visualization of CD34+ cells in BM, PB and mPB. **b**. Tissue source of each cell subset. **c**. Density plot of cells along the pseudotime in each tissue. **d**. Heatmap of expression of DEG modules along pseudotime. **e**. The top enriched GO terms of each module. **f-h**. UMAP visualization of the expression of DEG module genes, CXCR4 (**f**), AREG (**g**) and JUND (**h**). **i**. Heatmap of DEGs between tissues in HSCs.

We identified differentially expressed genes (DEGs) along pseudotime for better understanding of gene expression dynamics during hematopoiesis. These DEGs were classified into 5 modules, among which module2 and module1 are the most significant modules (**Fig 2d**). Module2 exhibits differences in HSC, MPP#1, MPP#2, LMPP#1, LMPP#2, GMP, MEP, CLP and EryP, while module1 exhibits differences mainly in MPP#1, MPP#2, LMPP#2, MEP, CLP, pro-B, EryP and MkP (**Fig 2d**). Therefore, module2 represents gene expression differences in early hematopoiesis, while module1 represents gene expression differences in hematopoiesis after HSCs. Module2 is significantly enriched in ribosome biogenesis and cytoplasmic translation (**Fig 2e**); while module1 is significantly enriched in leukocyte migration, positive regulation of cell adhesion, hemostasis and platelet activation (**Fig 2e**). These results indicate the main differences during hematopoiesis are genes associated with cell migration, cell adhesion, hemostasis and platelet activation, while the main difference in early hematopoiesis is ribosome biogenesis. The most significant DEGs along the pseudotime include *CXCR4*, *AREG* and *JUND* (**Fig 2f-h**)*. AREG*, a BM specific gene, regulated cell growth and cell proliferation. *JUND*, a PB/mPB specific gene, been proposed to protect cells from p53-dependent senescence and apoptosis.

We further identified DEGs of HSCs between BM and PB/mPB to study the differences of early hematopoietic subsets between tissues. Indeed, BM specific genes of HSC were enriched in cytoplasmic translation, ribonucleoprotein complex biogenesis, nucleic acid transport and RNA transport (**Supplementary Fig 2b**), which indicate that HSCs in BM prepares a lot RNA for the later differentiation since almost all the enriched GO terms are related to RNA process. PB/mPB specific genes were enriched in ATP metabolic process, aerobic respiration and mitochondrial respiratory chain assembly (**Supplementary Fig 2b**), indicating HSC in PB have increased energy usage since all the enriched GO terms are related to energy. The most significant BM-specific genes of HSC are ribosomal proteins (RPs) include *RPL27A*, *RPL13A*, *RPL21*, *RPS17* and *RPL31* (**Fig 2i**), among which *RPL31* has been reported to inhibit cell migration^22^. BM specific genes of MPP, LMPP, MEP and MkP are also enriched in RNA biogenesis and ribonucleoprotein biogenesis (**Supplementary Fig 2c-f**), similar to that of HSCs. Furthermore, HSCs have the highest HSC score while PB have high Ery score and Mk score (**Supplementary Fig 2g**), indicating HSPCs in BM have more differentiation potential and HSPCs in PB have more functional potential.

### Inference of potential mechanism of HSPCs anchoring and migration out of BM

HSC, pre-B and EryP have the lowest migration scores among all HSPC subsets (**Supplementary Fig 3a**), indicating these HSPC subsets tend to be static. The migration score of HSPC subsets from BM were the lowest among the three tissues (**Fig 3a**), potentially because BM HSPCs anchored on BM stromal while HSPCs in PB are moving. Analysis of cell-cell interaction among cell subsets showed BM stromal cell (BM_SC) have the strongest interaction with CD34+ cell subsets (**Fig 3b**). Cell-cell interaction signals with BM_SC showing significant differences among HSPC subsets include *CXCR4*-*CXCL12*, *FN1*-*CD44*, *APP*-*CD74*, *FN1-(ITGAV+ITGB1)* and *FN1*-*(ITGA5+ITGB1)* (**Fig 3c; Supplementary Fig 3b**), which belong to signals associated with cell adhesion and cell migration. *CXCR4*-*CXCL12* axis is one of the most variable interactions signaling pathway among HSPC subsets. Interaction analysis of *CXCR4*-*CXCL12* axis showed that BM_SC was the strongest signaling sender through *CXCL12*, and pre-B was the strongest signaling receiver through *CXCR4* among all the cell subsets in BM (**Fig 3d**). Indeed, pre-B expressed the highest level of *CXCR4* and BM_SC expressed the highest level of *CXCL12* among all cell subsets in BM (**Fig 3e, f**). Furthermore, expression of *CXCR4* in PB/mPB is much lower than that in BM, potentially indicates decreased expression of *CXCR4* leads to weaker *CXCR4*-*CXCL12* interaction thus facilitate HSPCs migration out of BM, consistent with reports that *CXCL12*-*CXCR4* axis played a crucial part in cell migration cross tissues^23–27^. Indeed, chemotaxis assay of mouse HSPCs showed that *Cxcl12* have pronounced efficient chemoattraction of HSPCs (**Fig 3g, h**). Interestingly, *CXCR4*^low^ HSPCs showed much quicker migration than *CXCR4*^high^ HSPCs (**Fig 3i, j**). Furthermore, BM endothelial cells strongly communicated with CD34+ cell subsets based on SELPLG signaling which facilitated cell adhesion of MPPs and LMPPs in BM (**Supplementary Fig 3c, d**).

**Fig 3.**
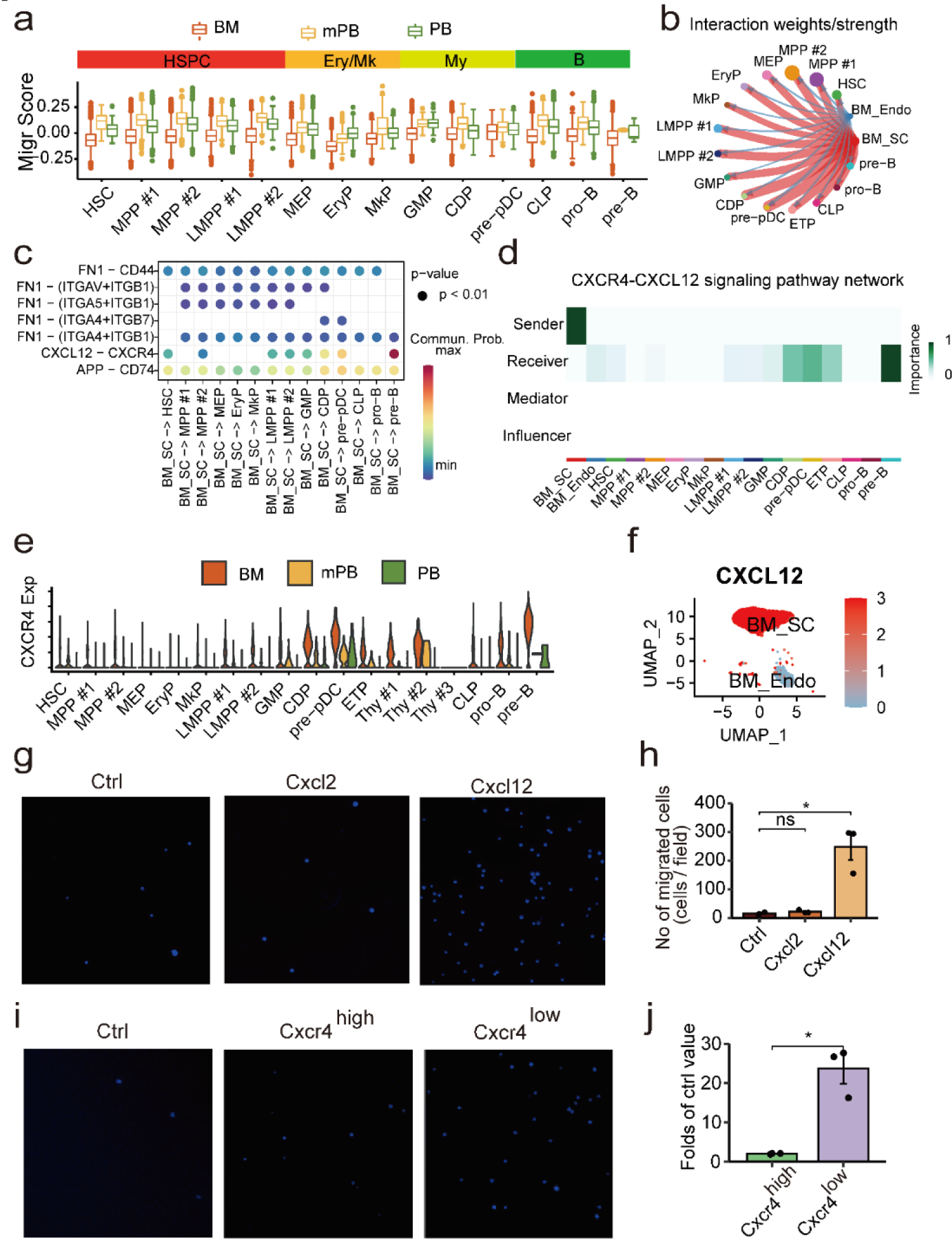
Inference of the potential mechanism of CD34+ cells remaining in or migration out of BM. **a**. Box plot of migration scores of each CD34+ cell subset in each tissue. **b**. Cell–cell interaction strength from BM stromal cell subsets to CD34+ cell subsets inferred by CellChat. The width and color of the line represent strength of cell-cell interaction and signaling source, respectively. Abbreviation: SC, stromal cells; Endo, endothelial cells. **c**. The signals of cell-cell interaction between HSPC subsets and BM_SC showing differences among HSPC subsets. **d**. Heatmap of *CXCL12*-*CXCR4* signaling between cell subsets in BM. **e**. Violin plot of the expression of *CXCR4* in each HSPC subset in each tissue. **f** UMAP visualization of the expression of CXCL12 in BM stromal cells. **g-h**. Chemotaxis assay of HSPCs showing image of DAPI staining (**g**) and number of migrated cells (**h**), with no chemokine (Ctrl), 100ng/ml *CXCL2* and 100ng/ml *CXCL12*. **i**. Image of DAPI staining of CXCR4^high^ HSPCs and CXCR4^low^ HSPCs, chemoattracted by 100ng/ml *CXCL12*. **j**. Fold change of the number of migrated cells in CXCR4^high^ HSPCs and CXCR4^low^ HSPCs compared with control.

### T cell compartment and change of cellular states cross tissues

Although the cell subsets in T lineages come from different tissues, they form a single connected entity on UMAP (**Fig 4a**). Each T progenitor cell subset is mainly composed of cells either from thymus or from BM/PB/mPB, except ETPs which consist of 52.9% of cells from thymus and 47.1% of cells from BM/PB/mPB (**Fig 4b**). Analysis of cell density along T lineage showed that HSPCs from BM dominated the earliest T lineage and their density decreased gradually along the lineage (**Fig 4c**). The cell density of HSPCs from PB/mPB peaked at MPP, while HSPCs from thymus were absent at early T lineage and dominated the T lineage after ETP. The decreasing curve of cell density of BM HSPCs and the rising curve of cell density of thymus HSPCs cross at ETP (**Fig 4c**), indicating that ETPs play an important role in the migration of HSPC from BM/PB to thymus. These results are consistent with reports that ETPs enter thymus for thymocyte differentiation^21,28,29^.

**Fig 4.**
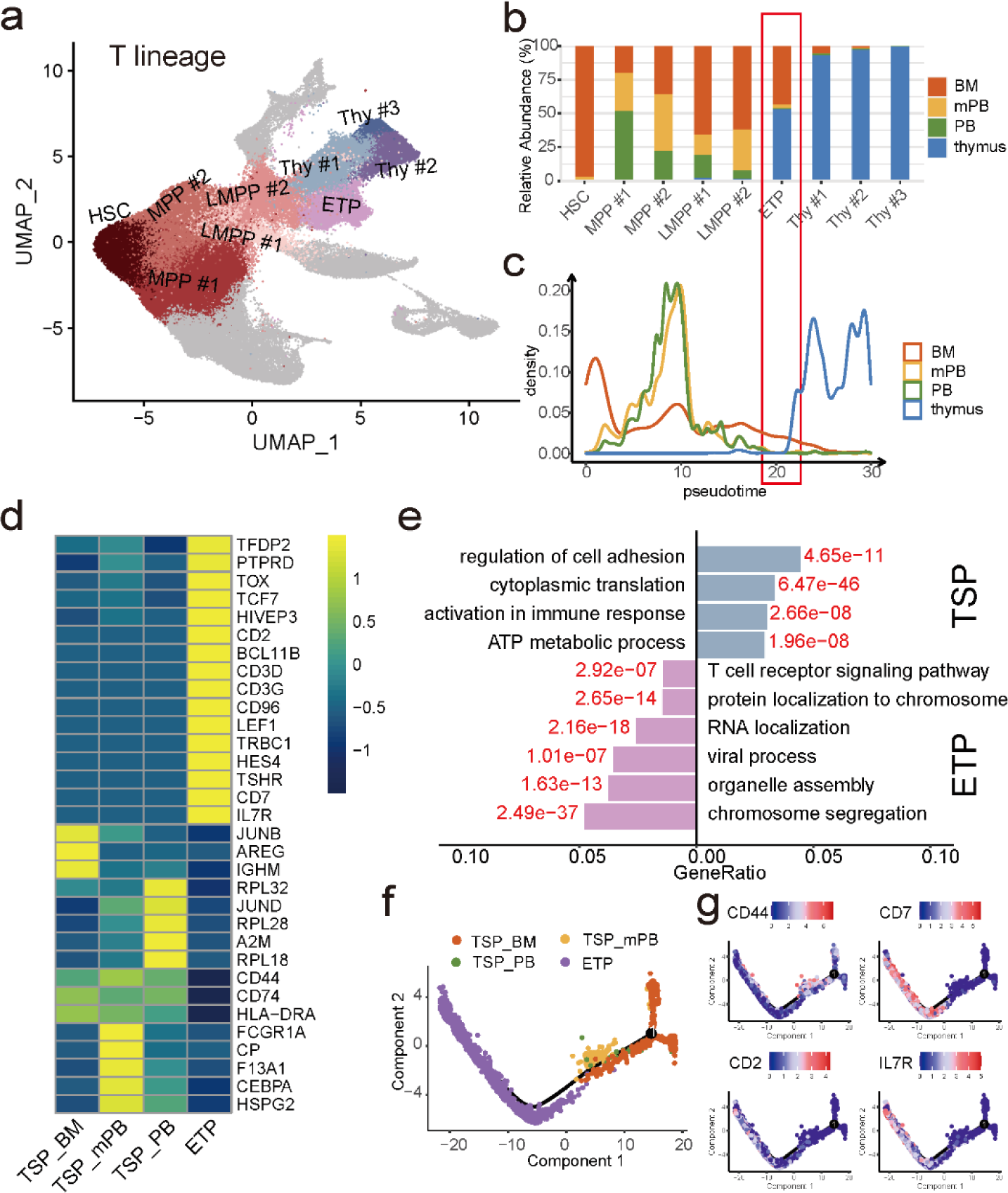
Early T cell compartment cross tissues and change of cellular states. **a.** UMAP visualization of CD34+ cells, with cells in T cell lineage being highlighted. **b.** Tissue source of each cell subset in T lineage. **c**. Density plot of cells along the T cell lineage in each tissue, with potentially ETP and TSP being highlighted. **d**. Heatmap of differentially expressed genes (DEGs) between ETP subsets. **e**. Enriched GO terms of DEGs between ETP and TSPs. **f**. Pseudotime analysis of cells from 4 ETP subsets indicates the developmental trajectory. **g**. Expression of indicated genes along the trajectory.

We focused on ETP subsets to better understand the migration of early T progenitors. ETPs in BM, mPB and PB were the progenitors for seeding in thymus, thus were called TSP_BM, TSP_mPB and TSP_PB, respectively. UMAP visualization showed that ETP subsets are different cell subsets (**Supplementary Fig 4a**). We identified 448 TSP specific genes and 1,012 ETP specific genes by comparing TSP subsets and ETP. The most pronounced TSP specific genes include ribosomal genes (*RPL4* and *RPL12*) and stem cell associated genes (*AREG*) (**Fig 4d**). The most pronounced ETP specific genes, including thymocyte determination genes (*CD7*, *CD3D*) (**Fig 4d**). ETP specific genes were enriched in T cell receptor signaling pathway while TSP specific genes were enriched in cell adhesion and immune response (**Fig 4e**).

TSP subsets highly expressed HSC specific genes, myeloid specific genes and B specific genes, while ETP expressed T lineage specific genes much higher than TSP subsets (**Supplementary Fig 4b-f**). Furthermore, TSP_BM specific genes were enriched in regulation of cell-cell adhesion and regulation of hematopoiesis, TSP_PB/TSP_mPB specific genes were enriched in positive regulation of cell adhesion and immune response (**Supplementary Fig 4g**). The differences between TSP subsets might be caused by differences of tissues microenvironment. Trajectory inference by Monocle 2^30,31^ showed that TSP_BM differentiated into TSP_PB/TSP_mPB, then differentiated into ETP (**Fig 4f**). The expression of *CD44* decreases along the trajectory (**Fig 4g**), consistent with recent studies that the expression of *CD44* decrease in ETP^18,32^. On the other hand, the expresses of *CD2*, *CD7*, *IL7R* increase along ETP trajectory (**Fig 4g**), consistent with recent studies that the expression of these genes increase in ETP^18,28^.

### Inferring the mechanism of TSPs homing to thymus

TSP_mPB and TSP_PB have higher migration scores than TSP_BM and ETP (**Fig 5a**), indicating the migration ability of TSP increases after moving from BM to PB and its migration ability decreased after homing to thymus. Analyses of ligand-receptor interaction from thymic stromal cell subsets to ETP subsets showed that thymic mesenchymal cells (thymic_Mesen) is the strongest signaling sender in thymus and is more likely to interact with TSP_mPB and TSP_PB (**Fig 5b**), potential indicating that thymic_Mesen and TSP_PB interaction play an important role in TSP_PB homing to thymus. The ligand-receptor interaction signaling between thymic_Mesen and ETP subsets showed two modes (**Fig 5c; Supplementary Fig 5a-i**). Mode#1 shows that interaction signaling with thymic_Mesen is enhanced along TSP_BM, TSP_PB and ETP, which include *CD99*-*CD99* and *PTN*-*NCL*. Mode#2 shows high interaction signaling with TSP_BM and ETP while low interaction with TSP_PB, which include *CXCL12*-*CXCR4*, *DLK1*-*NOTCH1*, *MIF*-*CD74*+*CXCR4* and *MIF*-*CD74*+*CD44* (**Fig 5c; Supplementary Fig 5a-i**). Expression level of genes in these interaction pairs in ETP subsets and thymic stromal subsets are essentially consistent with the two interaction modes (**Fig 5d**). In particular, the expressions of *CD99* and *NCL* increased along TSP_BM, TSP_PB and ETP (**Fig 5d**), consistent with interaction mode#1. Further qPCR analyses showed that expression level of *Cd99* increased along ETP trajectory (**Fig 5e**). The expressions level of *CXCR4* and *NOTCH1*, are high in TSP_BM and ETP while their expressions are low in TSP_PB (**Fig 5d**), consistent with interaction mode#2. qPCR and MACS analyses showed *Cxcr4* highly expressed in TSP_BM and ETP and lowly expressed in TSP_PB (**Fig 5f, g**). These results are consistent with a report that noncanonical *DLK1*-*NOTCH* signal potentially promoted ETP maturation^33^.

**Fig 5.**
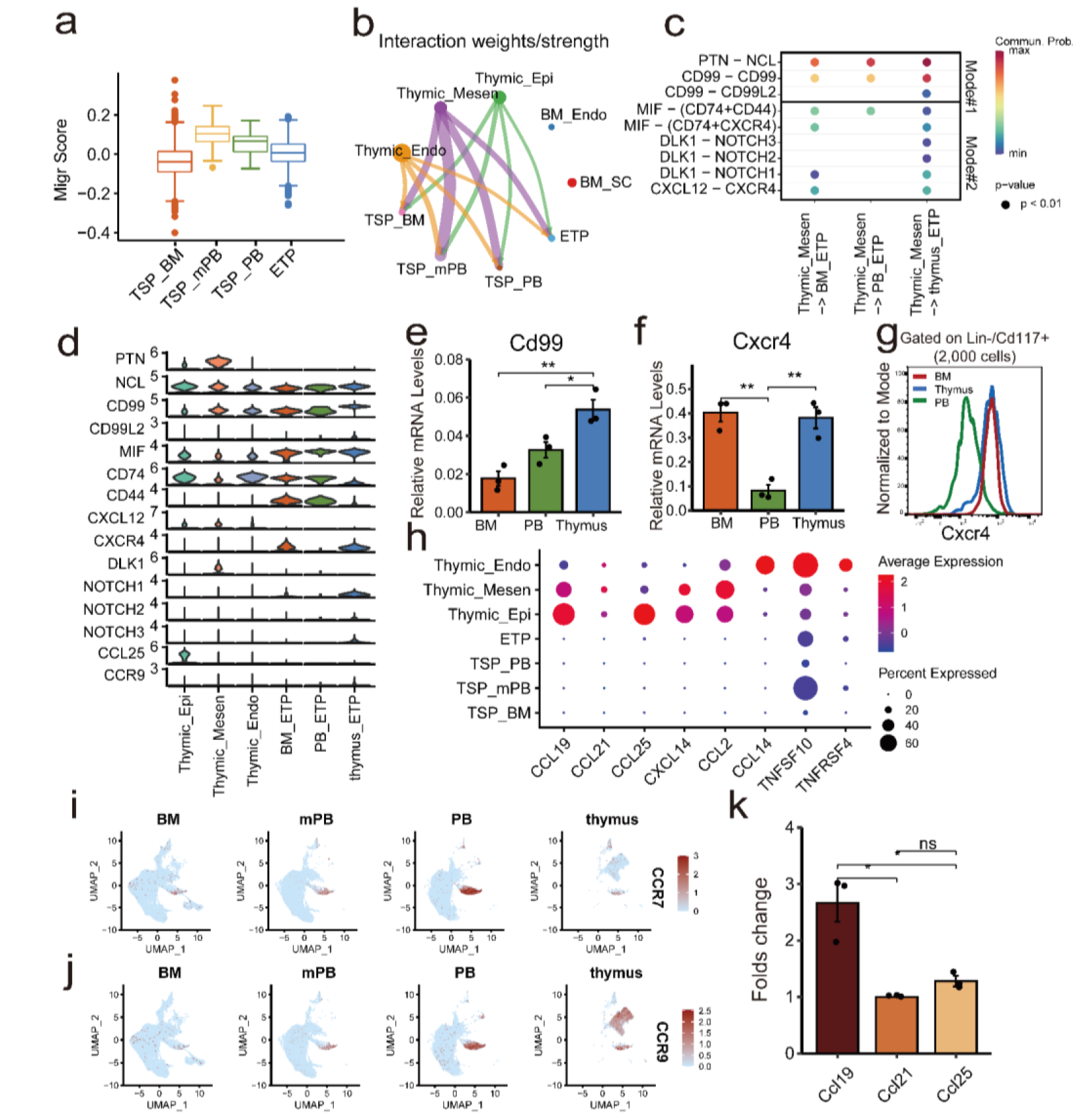
Inference of potential mechanism of ETPs homing to thymus. **a.** Boxplot of migration scores of ETPs in each tissue. Migration score was calculated by the accumulation of gene expression of migration related genes. **b**. Inferred cell–cell interaction strength among thymic epithelial cells (Thymic_Epi), thymic mesenchymal cells (Thymic_Mesen), thymic endothelial cells (Thymic_Endo) and the 4 ETP subsets. **c.** The significantly changed interaction signaling between ETP subsets and Thymic_Mesen. **d**. Expression of genes involved in interaction signaling in ETP subsets and thymic stromal cell subsets. **e-f**. Expression level of Cd99 (**e**) and Cxcr4 (**f**) in mouse BM, PB and thymus by qPCR. **g**. FACS analysis of *CXCXR4* in BM, PB and thymus. **h**. Dot plot of the expression level of cytokines in ETP subsets and thymic stromal cell subsets. **i-j**. Expression of *CCR7* (**i)** and *CCR9* **(j)** in the four tissues. **k**. Fold change of number of migrated cells chemoattracted by *CCL19*, *CCL21* and *CCL25* compared with control.

Although cell-cell interaction through ligand-receptor pair provides biological insight into ETP homing thymus, we are also interested in how chemokines affect ETP homing thymus. We found that thymic stromal cell subsets highly expressed chemokines such as *CCL19*, *CCL21*, *CCL25, CXCL14, CCL2*, *CCL14, TNFSF10* and *TNFSFR4* (**Fig 5h**). We found that HSPCs expressed *CCR7* and *CCR9* in PB and thymus (**Fig 5h-j**). Therefore, we assume that *CCL19*-*CCR7* and *CCL25*-*CCR9* might play important roles in ETP homing to thymus. Chemotaxis assay showed that *CCL19* attracted significantly more HSPCs than control (**Fig 5k; Supplementary Fig 5j**), consistent with reports that *CCL19* with its receptor *CCR7* regulates ETP homing thymus^34,35^.

### Abnormalities of feature and migration ability of HSPCs in MF patients

MF is a myeloproliferative neoplasm that disrupts the body’s normal production of blood cells. It is reported that MF has increased numbers of abnormal Mks and lead Mk-biased hematopoiesis^13^. We integrated 82,561 mPB CD34+ cells from 15 MF patients from Psaila et al.^13^ for further investigating the abnormalities of hematopoiesis. Projecting these HSPCs from MF patients on the reference healthy HSPCs atlas showed a hierarchical structure, which are essentially consistent with that of healthy reference (**Fig 6a**). However, the cell subset and population size in MF patients are different from that of healthy individuals (**Fig 6b**). In particular, MkP, EryP, LMPP#1, GMP and pre-pDC are mainly from MF patients while CLP, pro-B, CDP and MPP#1 are mainly from healthy individuals, consistent with the report that there are increased Mk lineage and decreased B lineage in MF^13^. Although there are a few HSCs in healthy mPB HSPCs, HSPCs from MF patients have a much higher fraction of HSCs, accounting for >50% of HSCs (**Fig 6b**), potentially indicating that the obstruction of hematopoiesis leads to the increase of HSC. HSC scores of HSPC subsets from MF patients were higher than their counterparts in healthy mPB (**Fig 6c**, **Supplementary Fig 6a, b**), potentially because of the increased stemness due to leukemogenesis. Furthermore, MF patients have increased Mk score and decreased B score than their healthy counterparts which indicated the raised Mk potential and reduces B potential in MF patients (**Fig 6d, 6e**).

**Fig 6.**
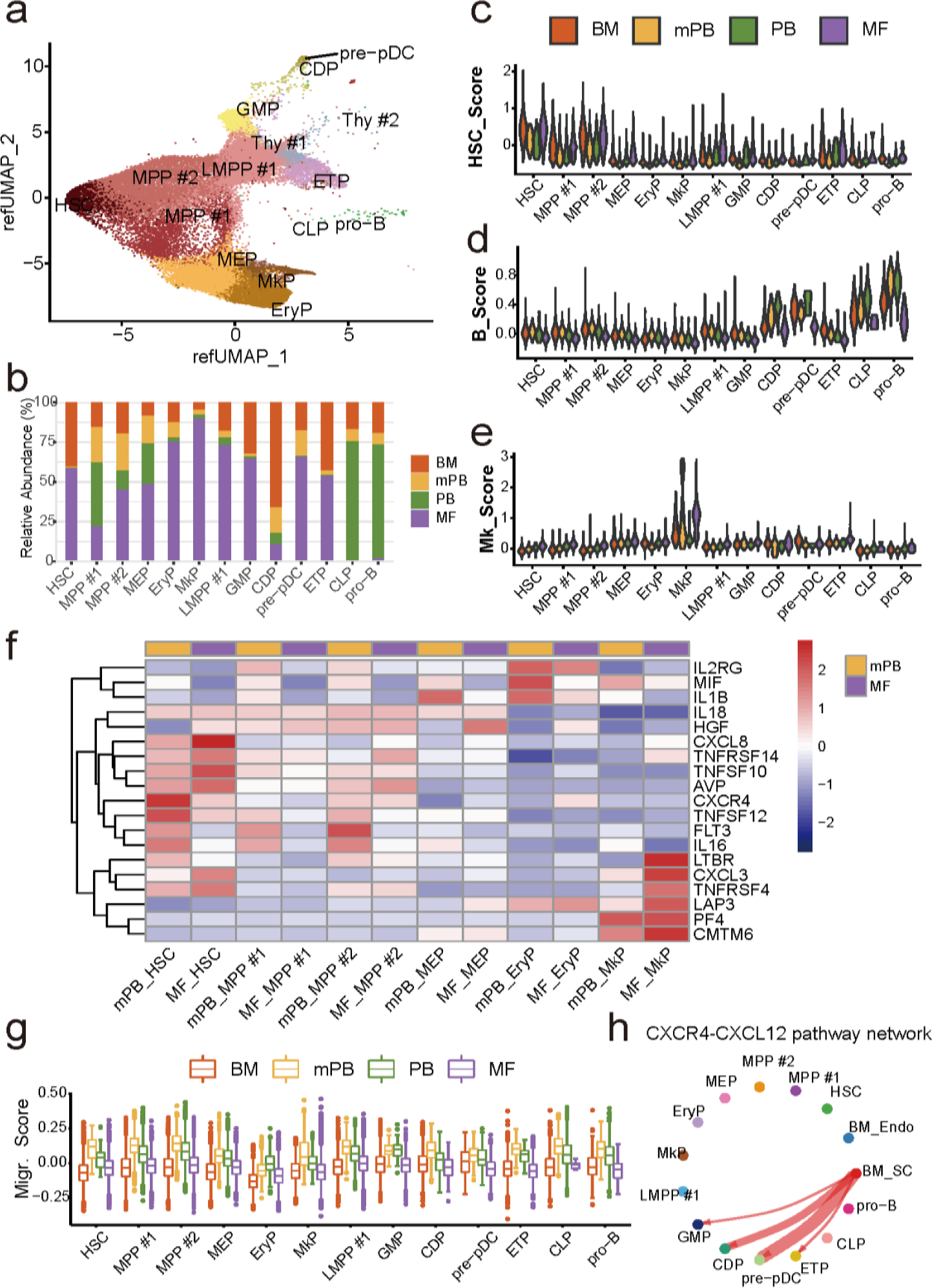
Abnormalities of population size, feature and migration ability of HSPC subsets in MF patients. **a.** CD34+ cell atlas of MF patients, constructed by projecting these cells on the reference healthy CD34+ cell atlas in Fig. 1b. **b**. Tissue or patient source of each cell subset, in which MkP, EryP, LMPP #1, GMP and pre-pDC were dominated by cells from MF patient, while almost absence from CLP, pro-B, GMP and MPP#1. **c-e**. HSC score **(c)** B score **(d)** and Mk score **(e)** of HSPC subsets in BM, PB, mPB and MF patient. **f**. Comparison of expression of chemokines and receptors in HSPC subsets in mPB and MF patient. **g**. Boxplot of migration score of HSPC subsets in BM, PB, mPB and MF patient. **h**. Cell communication network among BM_SC, BM_Endo and HSPC subsets in MF patients based on CXCL signaling pathway.

We identified 1,031 DEGs between MF patients and healthy individuals, in which the most significant enriched genes include genes involved in cytokines signaling and cell adhesion (**Fig 6f**). The cytokines signaling, such as *IL18*, *IL16*, *IL1B* and *IL2RG*, play important roles in immunomodulatory cytokine networks involved in host defense^36^. Interestingly, the migration scores of HSPC subsets in MF patients were much lower than that in their healthy counterparts (**Fig 6g**), reflecting that the decreased mobility of HSPCs in PB is an unreported cellular dysfunction in MF patients. MF patients have the weakest interaction between BM_SC and HSC based on *CXCL12*-*CXCR4* signaling (**Fig 6h**; **Supplementary Fig 6c-f**).

### A model of HSPC differentiation and migration

We proposed a model to describe HSPCs differentiation and migration from BM, to PB/mPB and thymus (**Fig 7**). In this model, HSCs in BM differentiated into various hematopoietic lineages with a hierarchical structure. Because HSPC subsets could freely migrate to PB from BM except HSC and B cell subsets in early B lineage, the inferred pseudotime of HSPCs in PB also showed a hierarchical hematopoiesis structure similar to that in BM. PB and mPB have very similar hierarchical structures and features. Among all HSPC subsets in thymus, we only found the counterparts of ETPs in BM and PB, indicating that ETPs were the subsets that migrated from BM to PB, then seeds in thymus for differentiating into various T cell subsets. In brief, the model of HSPCs homing to thymus was completely different from that from BM to PB.

**Fig 7.**
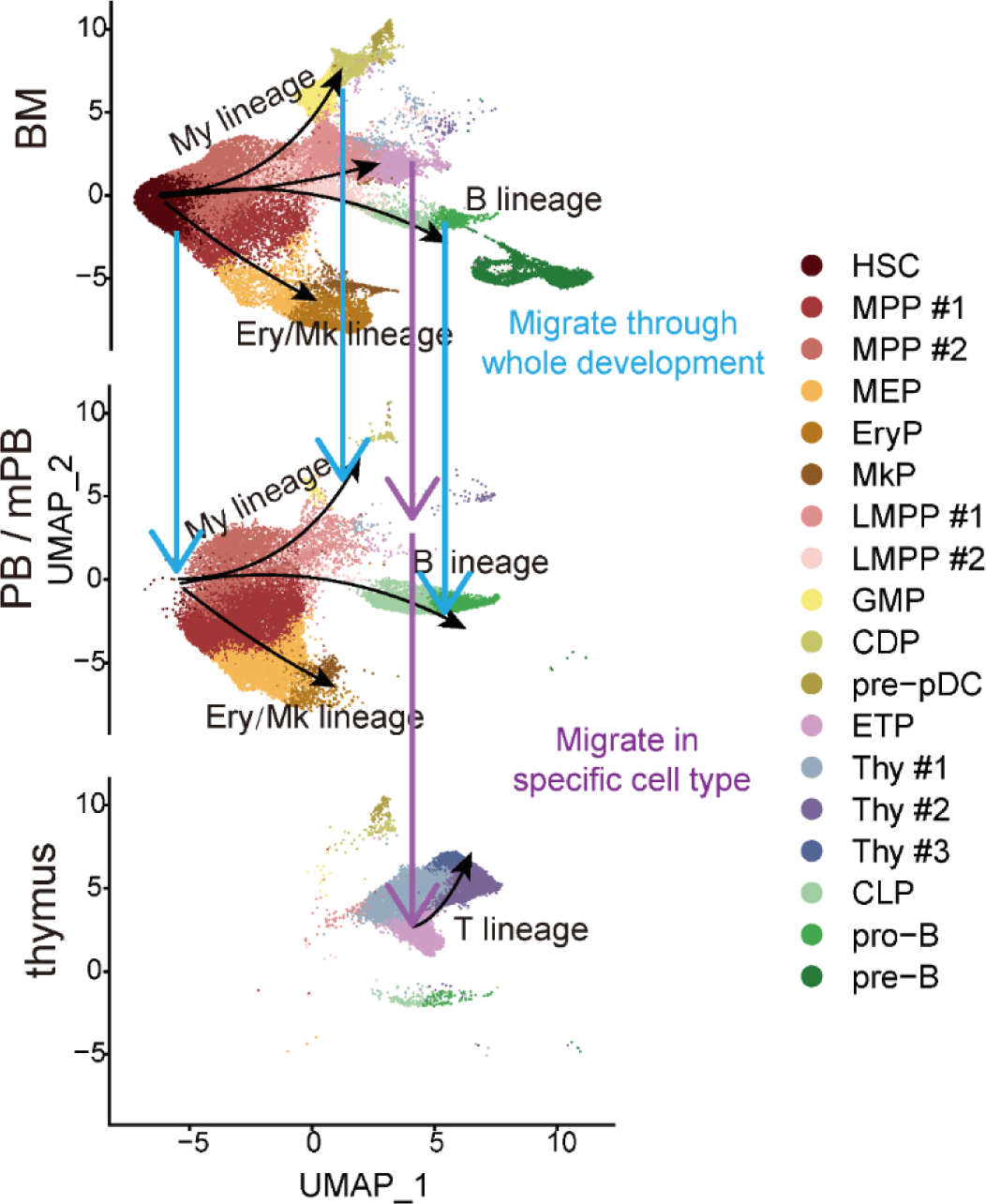
The model of CD34+ cells differentiation and migration cross BM, PB/mPB, and thymus. Supplementary Figures.

## Discussion

The scRNA-seq greatly expanded our understanding of early hematopoiesis in BM^1–3^. However, most studies only focused on hematopoiesis in one tissue and did not consider the multi-tissues coordination during process of hematopoiesis. In this study, the scRNA-seq data of CD34+ cells in BM, PB, mPB and thymus were integrated for constructing the cell atlas of HSPCs and inferring a relatively complete hematopoietic process. The inferred hematopoietic lineage of BM, PB and mPB are similar to each other, showing a continuous process with hierarchical structure. In particular, the hematopoietic lineage inferred with mPB CD34+ cells are very similar to that inferred with PB CD34+ cells. The inferred hematopoietic lineage in BM reflects a real hematopoietic process, while the continuous hematopoietic lineage in PB/mPB reflects that PB/mPB has complete HSPC spectrum or almost complete HSPC subsets which are mainly migrated from BM. In this way, although the inferred hematopoietic lineages in BM, PB and mPB were similar, the reasons for the formation of these similar hierarchical hematopoietic lineage are quite different. The HSPCs in thymus have several different subsets and mainly belong to T lineage. Among all HSPC subsets in thymus, we only found the counterparts of ETPs in BM and PB, indicating that ETPs were the subsets that migrated from BM to PB, then seeding in thymus. We found *CCL19*-*CCR7* might play a role in ETP homing to thymus. In brief, the migration of HSPCs from BM to PB is almost a full spectrum migration, while HSPCs homing to thymus is a small HSPC subsets seeding process, representing two quite different models of HSPCs migration.

Although the inferred hematopoietic lineage of BM, PB and mPB are similar to each other, each HSPC subset has a lot DEGs between tissues. The same cell subset has different expression profiles in different tissues, which is mainly caused by different local microenvironments in different tissues. We calculated the DEGs along the pseudotime which showed several modules. During the early hematopoiesis, BM specific genes of HSC were enriched in ribosome biogenesis, cytoplasmic translation, nucleic acid transport and RNA transport, and highly expressed ribosomal proteins (RPs) include *RPL27A*, *RPL13A*, *RPL21*, *RPS17* and *RPL31*, which indicates that HSCs in BM prepare a lot RNA for the later differentiation. PB/mPB specific genes of HSC were enriched in ATP metabolic process and aerobic respiration, indicating HSC in PB have increased energy usage. The most significant DEG module between tissues is module1, which significantly enriched in leukocyte migration, positive regulation of cell adhesion and hemostasis, indicating the main difference during hematopoiesis are genes associated with cell migration, cell adhesion and hemostasis. Although mPB CD34+ cells were used for BM transplants, they are similar to PB CD34+ cells instead of BM CD34+ cells. The similarity between mPB HSPCs and PB HSPCs may be caused by the same microenvironment. Compared with PB HSPCs, mPB HSPCs expressed more stemness genes and have increased mobility, potentially caused by the premature migration out of BM. For the ETP subsets, TSP subsets highly expressed HSC specific genes, myeloid specific genes and B specific genes, while ETP expressed T lineage specific genes much higher than that TSP subsets.

We explored the potential mechanism of HSPCs migrating between tissues, focusing on the migration of HPSC from BM to PB and HPSC homing to thymus. We found that the interaction between BM_SCs and HSPCs through CXCL12-CXCR4 axis plays an important role in the retention of HSPCs in BM. In particular, pre-B has the highest expression of *CXCR4* among all HSPC subsets and almost exclusively locates in BM. The HSPC subsets in PB/mPB have much higher migration ability than their counterpart in BM and thymus. HSPCs homing and seeding in thymus is also an important event in hematopoiesis^29^. We showed that ETP is the only HSPC subset that could be found in thymus, BM, PB and mPB, indicating ETPs in thymus may derivate from ETPs in BM, namely TSP_BM. The expression of *NCL* and *CD99* increase along the trajectory of ETP subsets, as well as the re-expression *CXCR4* and *NOTCH1* in ETP, may facilitate ETPs homing to thymus. Although cell-cell interaction through ligand-receptor pair provides a lot biological insight into ETP homing to thymus, we identified chemokine factors such as *CCL19*-*CCR7* might play a role in ETP homing to thymus. Although CCR9 has been reported to play an important role in ETP homing to thymus^37–39^,we did not find chemoattraction of *CCL25* played important role in ETP homing to thymus. Furthermore, we found the migration score of HSPC subsets in MF patients is much lower, reflecting that the decreased mobility of HSPCs in PB is an unreported cellular dysfunction in MF patients.

In conclusion, we constructed cell atlas of HSPCs and inferred a relatively complete hematopoietic lineage. We analyzed differentiation of HSPCs in BM, migration from BM to PB, then homing to thymus at single cell transcriptome resolution, which could be used as reference for studies of hematological diseases and might provide new insights for clinical applications.

## Methods

### Single cell RNA-seq data and sample information

In order to systematically investigate the cross-tissue differentiation and migration of HSPCs, we obtained the scRNA-seq data of human CD34+ cells from our previous studies^4,40^ and public data^13,15,17,18^. In brief, samples of these scRNA-seq data include bone marrow CD34+ cells from Qin *et al.*^4^ and Hua *et al.*^15^, peripheral blood (PB) CD34+ CD19-cells from Mende *et al.*^17^, mobilized PB (mPB) CD34+ Lin-cells from Psaila *et al*.^13^, and thymus CD34+ Lin-cells from Le *et al.*^18^. The scRNA-seq data of HSPCs from healthy individuals include 5 BM samples, 4 mPB samples, 4 PB samples, and 3 thymus samples. In total, we obtained 90,598 single cell transcriptomes of CD34+ cells in four tissues to investigate HSPCs differentiation and migration.

We further obtained scRNA-seq data of BM stromal cells from Li *et al.*^41^ for investigating the mechanisms of HSPCs migration out of BM. The scRNA-seq of thymic stromal cells were obtained from Bautista *et al.*^42^ for investigating the HSPCs seeding to thymus. In addition, we obtained mPB CD34+Lin-cells in 15 MF patients from Psaila *et al.*^13^ to investigate the abnormalities of HSPCs migration and differentiation.

### Mice samples and isolation of bone marrow c-Kit+ cells

The 8-12 weeks old C57BL/6J wild type mice were used for collecting of HSPCs. All mice experiments were approved by the Institutional Animal Care and Use Committee of Southern University of Science and Technology. The bone marrow lineage-(Lin-) and c-Kit/Cd117+ cells were isolated from wild type mice following our previous study^43^. Briefly, femurs and tibia from C57BL/6J mice were obtained and the BM cells were flushed out with 1×PBS supplemented with 0.1% BSA and 1 mM EDTA. The cell suspension was pass through a Falcon^®^ 40μm strainer (catalog no.352340, Corining). Red blood cells (RBC) were lysed using Red Blood Cell Lysis Solution (catalog no. 130-094-183, Miltenyi BIotec) according to the manufacturer’s instructions. Lineage cells were depleted using Mouse Lineage Cell Depletion Kit (catalog no. 130-110-470, Miltenyi BIotec) and pass through LD depletion columns (catalog no. 130-042-901, Miltenyi BIotec) according to the manufacturer’s instructions. The Cd117+ BM cells were further enriched using mouse Cd117 Selection Kit (catalog no. 18757, STEMCELL) according to the manufacturer’s instructions. The Lin-Cd117+ BM cells were kept on ice before further use.

### Pre-processing of scRNA-seq data

The fastq data were pre-processed similar to our previous studies^4,44^. Briefly, reads were aligned to human reference genome GRCh38 and raw matrix were generated using Cell Ranger 7.0.1^12^. The matrices were loaded into Seurat 4.1.2^19^ for further analysis. The cells with <1000 unique molecular identifier (UMI) counts, <200 number of detected genes or >10% percentage of mitochondria genes >10% were filter out. The scRNA-seq data of each tissue (i.e., BM, PB, mPB and thymus) were integrated using Canonical Correlation Analysis (CCA)^19^.

### Data integration, dimension reduction, clustering and annotation

Seurat ^19^ was used for data normalization, data integration, dimension reduction and cell clustering. The processed data from the four tissues were integrated together using CCA with parameters of 12 anchors and 20 dims. Dimension reduction and visualization of scRNA-seq were performed using Uniform Manifold Approximation and Projection (UMAP)^45^. Cells were clustered using k-nearest neighbor (KNN) graph with 0.95 resolution and 23 PCs. Genes exhibiting significantly higher expression within the investigated cluster compared with the remaining cells were called cluster-specific genes. Each cell cluster was annotated according to well-established cell type specific genes^4,18,46^ and cluster specific genes. Small clusters that expressed marker genes from two different cell types were considered as doublets and were excluded.

### Trajectory inference and hematopoietic lineages

Monocle 3 v1.2.9^30,31,45,47^ was used to infer the hematopoietic trajectory. Monocle 3 uses UMAP to embed cells in low dimensional space thus we could easily visualize the cellular relationship in well-established cell clusters. Monocle 3 uses principal graph embedding algorithms to learn a trajectory that fits the cells’ UMAP coordinates. We selected HSC as the start of hematopoietic lineages according to previous studies^4,15^. We identified 4 main hematopoietic lineages, namely Ery/Mk lineage, My lineage, B lineage and T lineages.

### Differentially expressed genes (DEGs) and gene oncology (GO) analysis

DEGs between two cell subsets or DEGs between two tissues of a cell subset were identified using Wilcoxon Rank Sum test. We only kept the DEGs that detected in at least 1% of the cells and their expression differences between the two tissues were at least 1.5-fold. DEGs along the trajectory were calculated using function *graph_test* which was based on the low dimensional embedding and the principal graph. GO analysis of DEGs was performed by *compareCluster* in clusterProfiler^48^. Redundant GO items were removed by setting 0.4 cutoff and the GO terms were ranked by adjusted p-value.

### Calculation of migration score and lineage score

Migration score and lineage score was calculated by averaging the expressions of migration related genes and lineage specific genes in each cell subset, respectively. Migration related genes were obtained from “GO:0030335 positive regulation of cell migration” in Quick GO (http://www.ebi.ac.uk/QuickGO)^49^. After removing the unexpressed genes in our data, we used *AddModuleScore* to calculate migration score. The lineage score of each cell was calculated using *AddModuleScore* based on the top 50 lineage specific genes of each lineage. The HSC score was calculated based on the top 50 HSC specific genes that identified by Laurenti *et al.*^50^. The Ery score and Mk score was calculated based on the 50 most significant EryP specific genes and the 50 most significant MkP specific genes in our data, respectively. The My score was calculated based on the 25 most significant CDP specific genes and the 25 most significant pre-pDC specific genes. Similarly, B score was based on pro-B and pre-B, while T score was based on Thy #1, Thy #2 and Thy #3.

### Analysis of cell-cell communications

Cell-cell communication signatures and communication strengths between cell subsets were inferred by using CellChat v1.6.1^51^. CellChat calculated communication probabilities of all pairs of ligands and receptors in each signaling pathways among cell subsets. We first calculated the overall interactions between stromal cell subsets and CD34+ cell subsets in BM or in thymus. Then we identified the most significantly cell-cell interaction signaling between two cell subsets. The significantly changed pathways between two cell subsets were calculated by rankNet function. The communication distance of a pathway is calculated by rankSimilarity function. Interaction weight between cell subsets is drawn by netVisual_circle function. Interaction signaling pathways in dot plot is drawn by netVisual_bubble function. Signaling communication probabilities in heatmap is drawn by netAnalysis_signalingRole_network function.

### ETP lineage scores along trajectory

ETP subsets were analyzed by UMAP with 25 PCs and their pseudotime was inferred by monocle 2 v2.24.2^30,31^. Lineage scores of ETP were drew in scatter plot with the smoothing method “gam” and 0.95 level of confidence interval by ggplot2.

### Chemotaxis Assay

The Lin-c-kit+ cells isolated from BM were used for investigation of chemoattraction of cytokines and cell migration. 5×10^6^ of Lin-Cd117+ cells were resuspended in 100μL RPMI1640 complete medium and seeded in the 24-well transwell plates with 5μm membrane pore size (catalog. 3421, Corning). 500μL medium alone or with 100ng/mL CXCL2 (catalog no. 250-15, PeproTech), CXCL12 (catalog no. 250-15, PeproTech), CCL5 (catalog no. 250-15, PeproTech), CCL19 (catalog no. 250-15, PeproTech), CCL21 (catalog no. 250-15, PeproTech) or CCL25 (catalog no. 481-TK-025/CF, R&D) was added to the bottom wells. Cells were incubated at 37℃ for 2 hours in a humidified incubator supplied with 5% CO_2_. The top wells were removed after 2 hours’ incubation and the migrated cells were stained with Hoechst 33342 (catalog. C1025, Beyotime) for 10 min.

### Comparison of Cxcr4 levels of c-Kit+ cells from BM, PB and thymus

The 8-12 weeks old C57BL/6J mice were anesthetized by intraperitoneal injection of 2.5% Avertin at 250mg/10g. PB was collected by intracardiac aspiration after euthanizing mice, followed by RBC lysis. BM cells were flushed out from the tibia and femurs and the lysed with RBC lysis buffer. Thymus was collected and gently dissociated using the end of the plunger of syringe. All the cells were pass through a Falcon^®^ 40μm strainer, and stained with Biotin anti-mouse Lineage Panel (catalog no. 133307, Biolegend), followed with Brilliant Violet 510™ Streptavidin according to the manufacture’s instruction. The lineage-labeled cells were further staining with anti-*Cd117* (catalog no.161503, Biolegend) and anti-*Cxcr4* (catalog no. 551967, BD Biosciences). The c-Kit+ cells were sorted out for RT-qPCR validation.

### RT-qPCR assay

Gene expression levels were quantified using RT-qPCR assay. Approximately 200,000 cells were collected for RNA extraction. Total RNA was extracted using TRIzol (catalog no. 15596026, Invitrogen) according to the manufacturer’s instructions. 1 μg of total RNA was used to synthesize cDNA using the ReverTra Ace qPCR Kit (catalog no. FSQ-101, TOYOBO) following to the manufacturer’s instructions. For RT-qPCR, the SYBR Green Realtime PCR master mix (catalog no. QPK-201, TOYOBO) in a final volume of 20μl. The PCR reactions were performed on an ABI StepOne Plus PCR system. The cycling conditions were as following: denaturation at 95 °C for 60s, followed by 40 cycles of denaturation at 95°C for 60s, annealing at 60°C for 15s, and extension at 72°C for 45s. Expression level of genes were normalized by the expression level of b2-microglobulin (B2M), a house-keeping gene. All primers were listed in the **Supplementary Table 2**.

### Statistical analysis

For all statistical analyses and data visualizations, we employed the programming languages R and Python. To identify the DEGs between two cell clusters, we employed the Wilcoxon Rank Sum test. Bonferroni correction was applied to conduct multiple tests.

## Acknowledgements

This study was supported by National Key R&D Program of China (2021YFF1200900, 2021YFA0909300), National Natural Science Foundation of China (32170646), Guangdong Basic and Applied Basic Research Foundation to N. H. (2023A1515011908), Shenzhen Science and Technology Program (JCYJ20220818100401003, KQTD20180411143432337) and open project of BGI-Shenzhen. This publication is part of the Human Cell Atlas – www.humancellatlas.org/publications/ [humancellatlas.org]. We thank all members from the Jin lab for the helpful discussion. We acknowledge the assistance of Core Research Facilities of SUSTech. We thank Xibin Lu for the excellent support of FACS. The computational work was supported by Center for Computational Science and Engineering at SUSTech.

## Author Contributions

W.J. conceived the project. H.L. conducted experiments with help from N.H., X.W. and G.C.. Y.Y. analyzed the data. W.J., X.Z. and Y.S. supervised the project. Y.Y. and W.J. prepared the manuscript, with all authors’ contribution.

## Supplementary Figures

**Supplementary Fig1.**
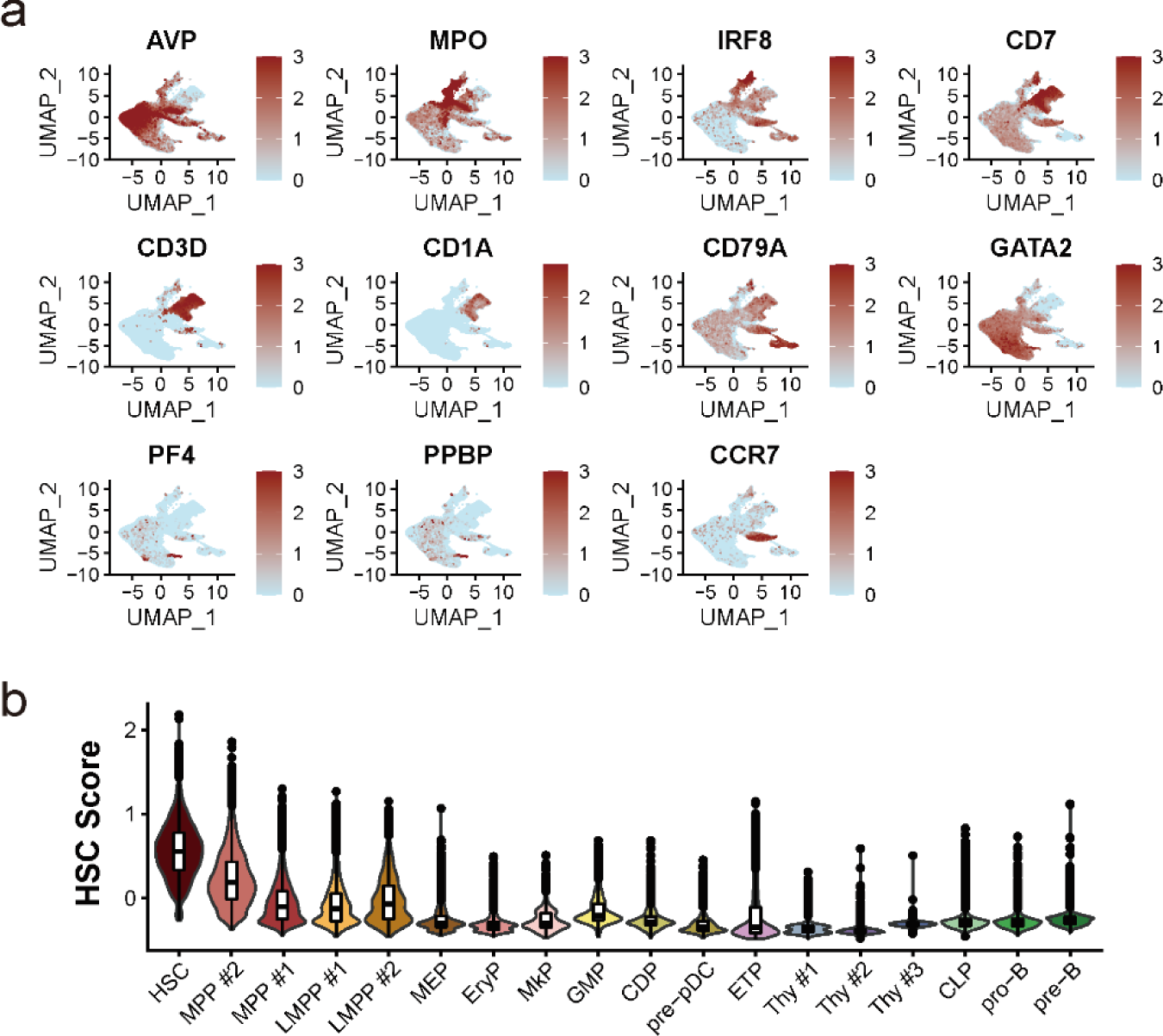
Cell subset specific genes and HSC scores. **a.** UMAP visualization of cell subset specific genes. **b**. Decreasing HSC score along pseudotime.

**Supplementary Fig2.**
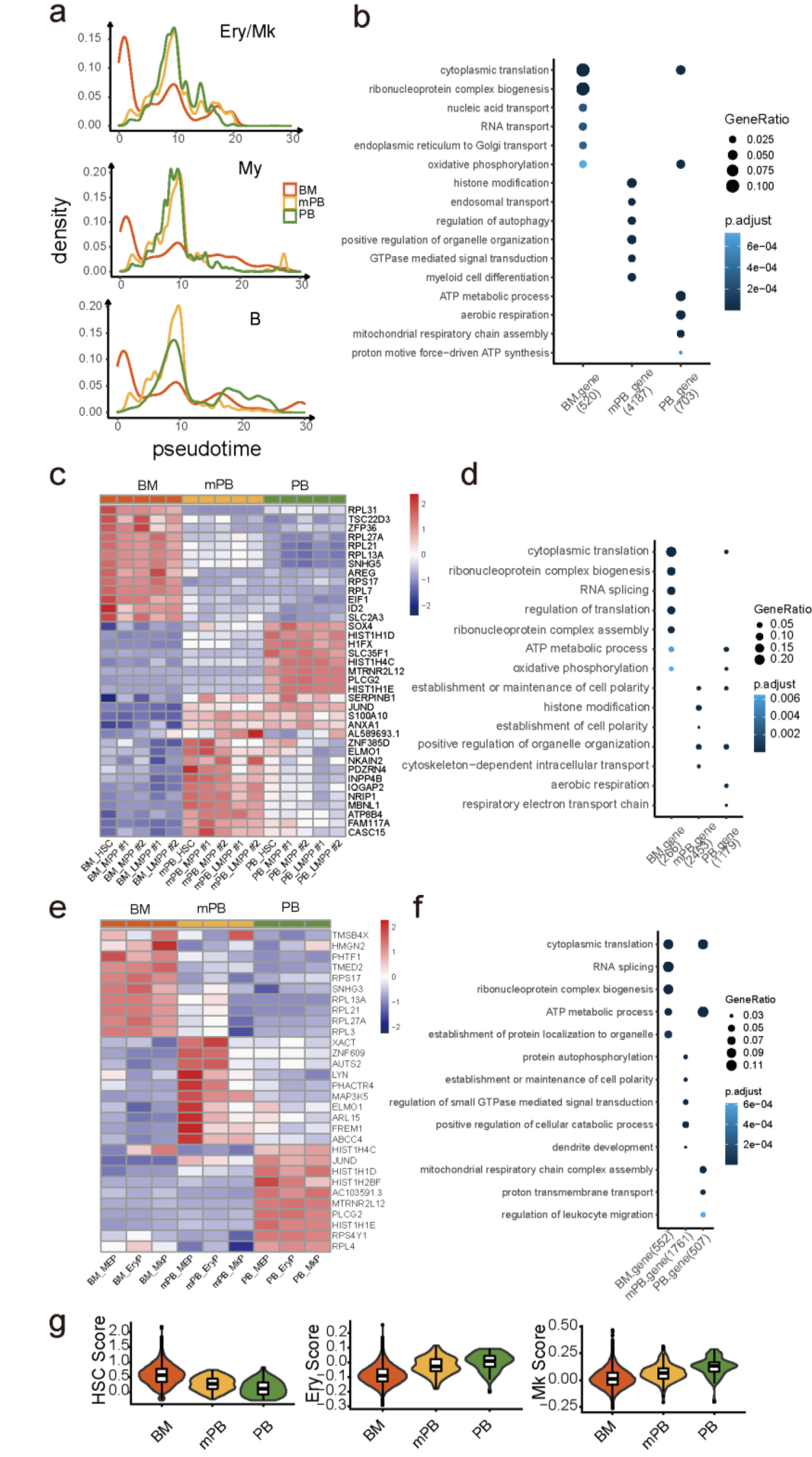
Cell density along lineage and tissue specific gene. **a**. Density plot of cells along in Ery/Mk lineage, My lineage and B lineage. **b**. GO enrichment of tissue specific genes in HSCs. **c**. Heatmap of top tissue specific genes in early hematopoietic HSPC subsets, namely HSCs, MPPs and LMPPs. **d**. GO enrichment of tissue specific genes of early hematopoietic HSPC subsets. **e**. Heatmap of top tissue specific genes in Ery/Mk lineages. **f**. GO enrichment of tissue specific genes of Ery/Mk lineages. **g**. Stemness score, Ery score, Mk score and My score of HSC cells in each tissue.

**Supplementary Fig3.**
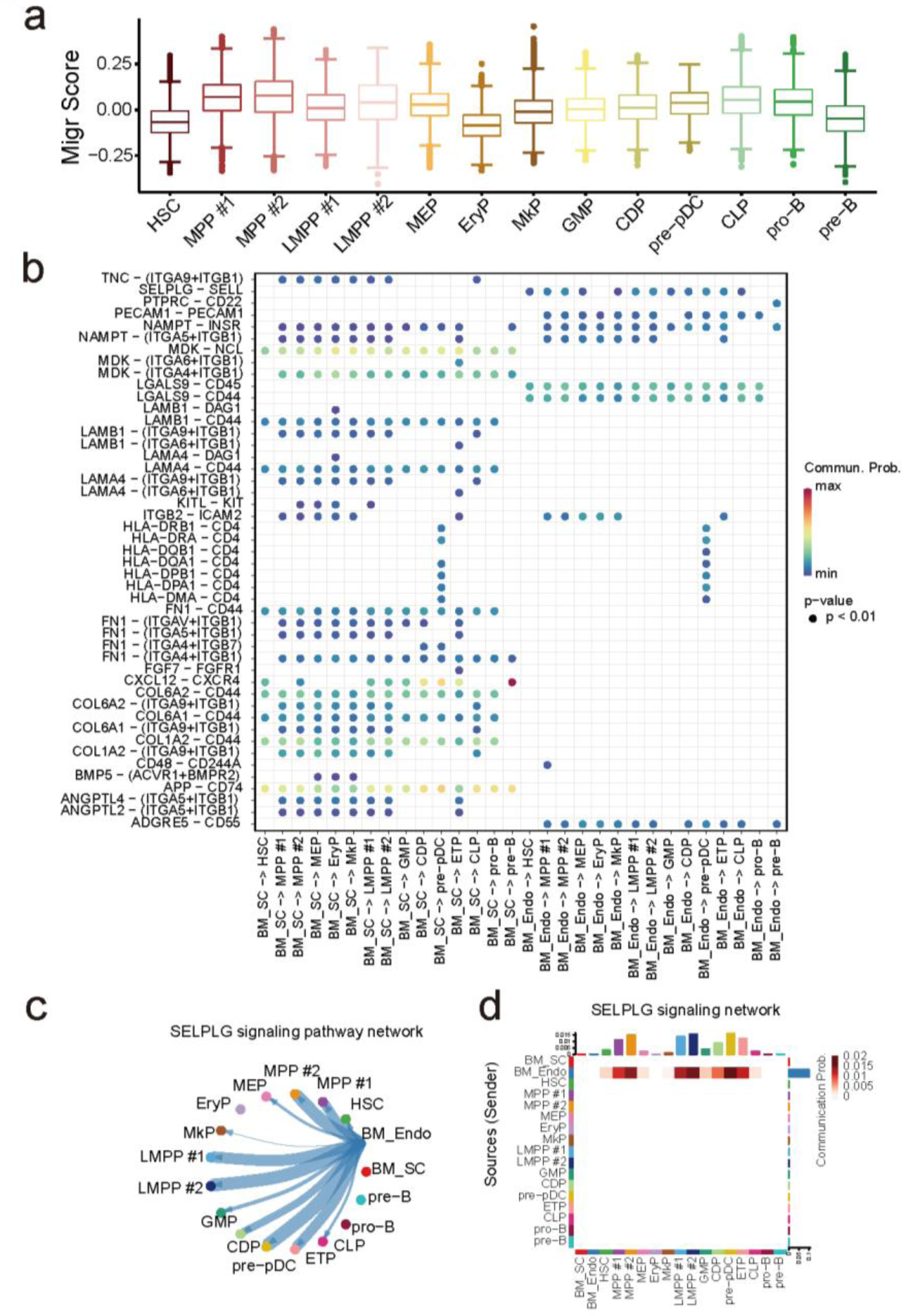
Comparison of migration score and interactions between BM stromal cell subsets to CD34+ cell subsets. **a.** Boxplot of migration score in each HSPC subset. **b**. Interaction pairs from BM stromal cell subsets to CD34+ cell subsets. **c-d**. Cell communication network **(c)** and heatmap of cell communication **(d)** from BM_Endo to CD34+ cell subsets based on SELPLG signaling.

**Supplementary Fig4.**
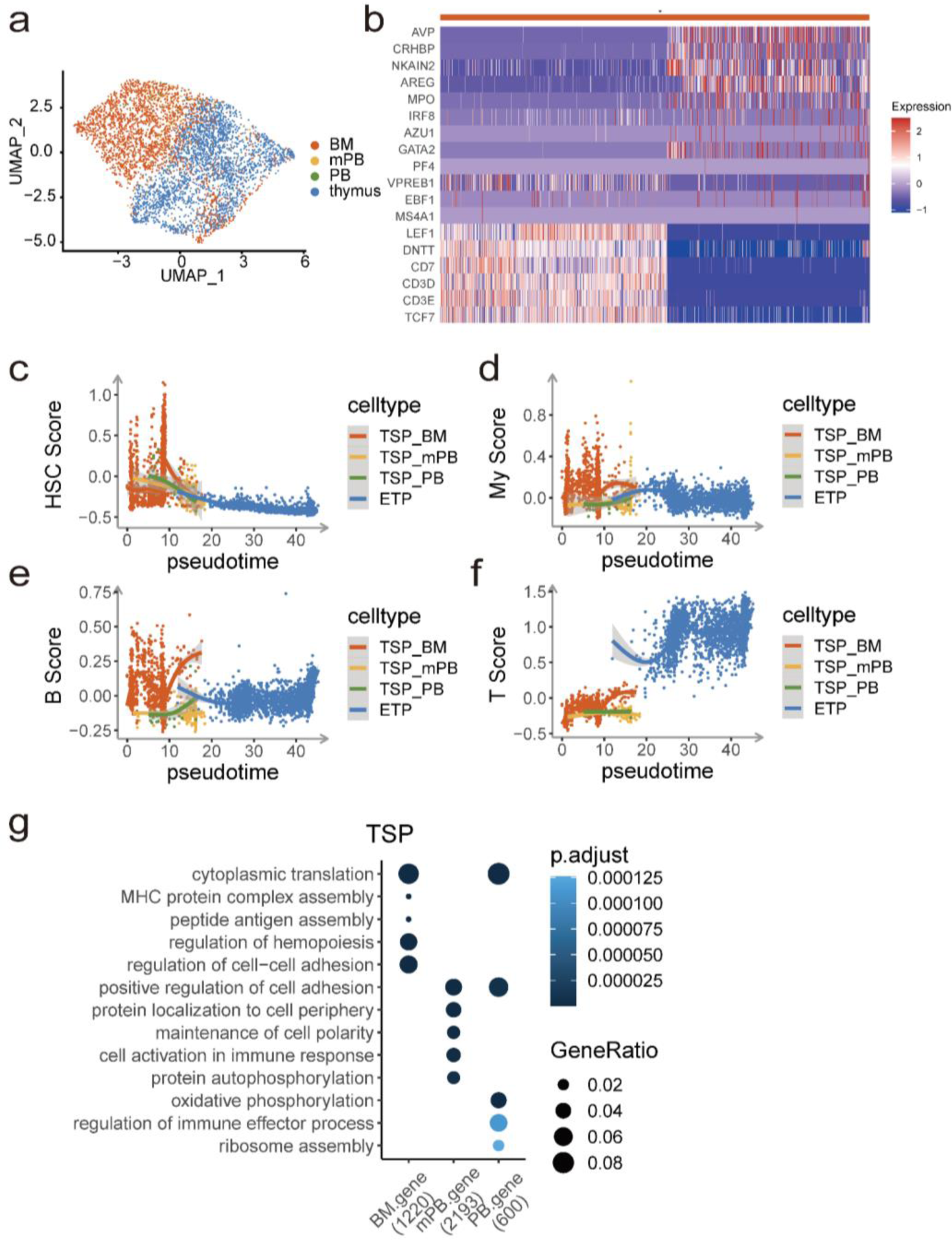
Comparison of ETP subsets for lineage potential and GO terms. **a.** UMAP visualization of ETP subsets, colored by inferred cell subset. **b**. Expression of lineage specific genes in ETP subsets. **c-f**. Stemness score **(c)**, My score **(d)**, B score **(e)** and T score **(f)** of ETP subsets along pseudotime. **g**. GO enrichment of tissue specific genes in TSP subsets.

**Supplementary Fig5.**
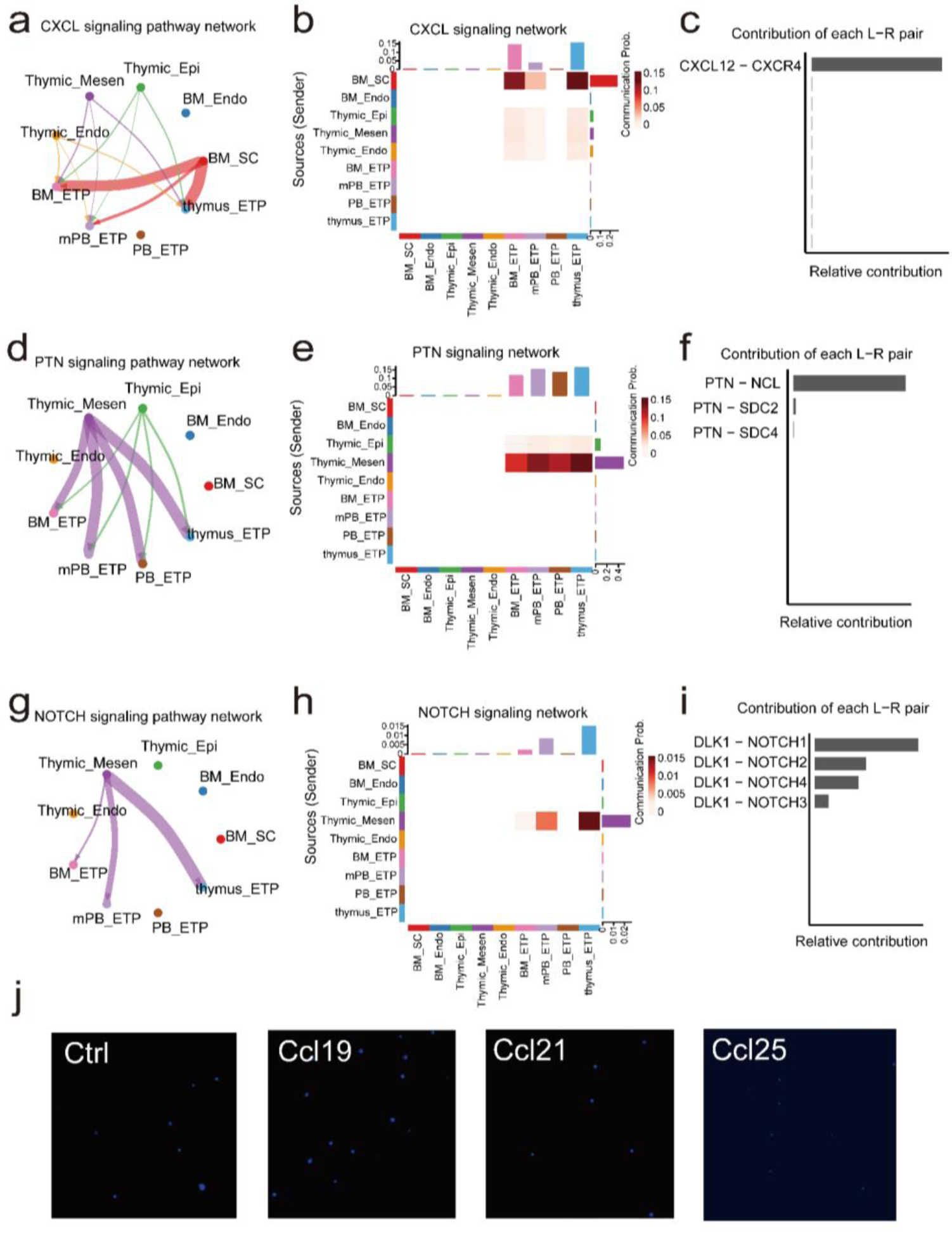
Potential cell signals from BM stromal cell subsets and thymic stromal cell subsets to ETP subsets. **a-c**. Cell-cell communication network (**a**), heatmap (**b**) and contribution of ligand-receptor pairs (**c**) from BM/thymic stromal cell subsets to ETP subsets based on CXCL signaling. **d-f**. Cell-cell communication network (**d**), heatmap (**e**) and contribution of ligand-receptor pairs (**f**) from BM/thymic stromal cell subsets to ETP subsets based on PTN signaling. **g-i**. Cell-cell communication network (**g**), heatmap (**h**) and contribution of ligand-receptor pairs (**i**) from BM/thymic stromal cell subsets to ETP subsets based on NOTCH signaling. **j**. DAPI staining of migrated cells with no chemokine (Ctrl), chemoattracted by 100ng/ml CCL19, CCL21 and CCL25.

**Supplementary Fig6.**
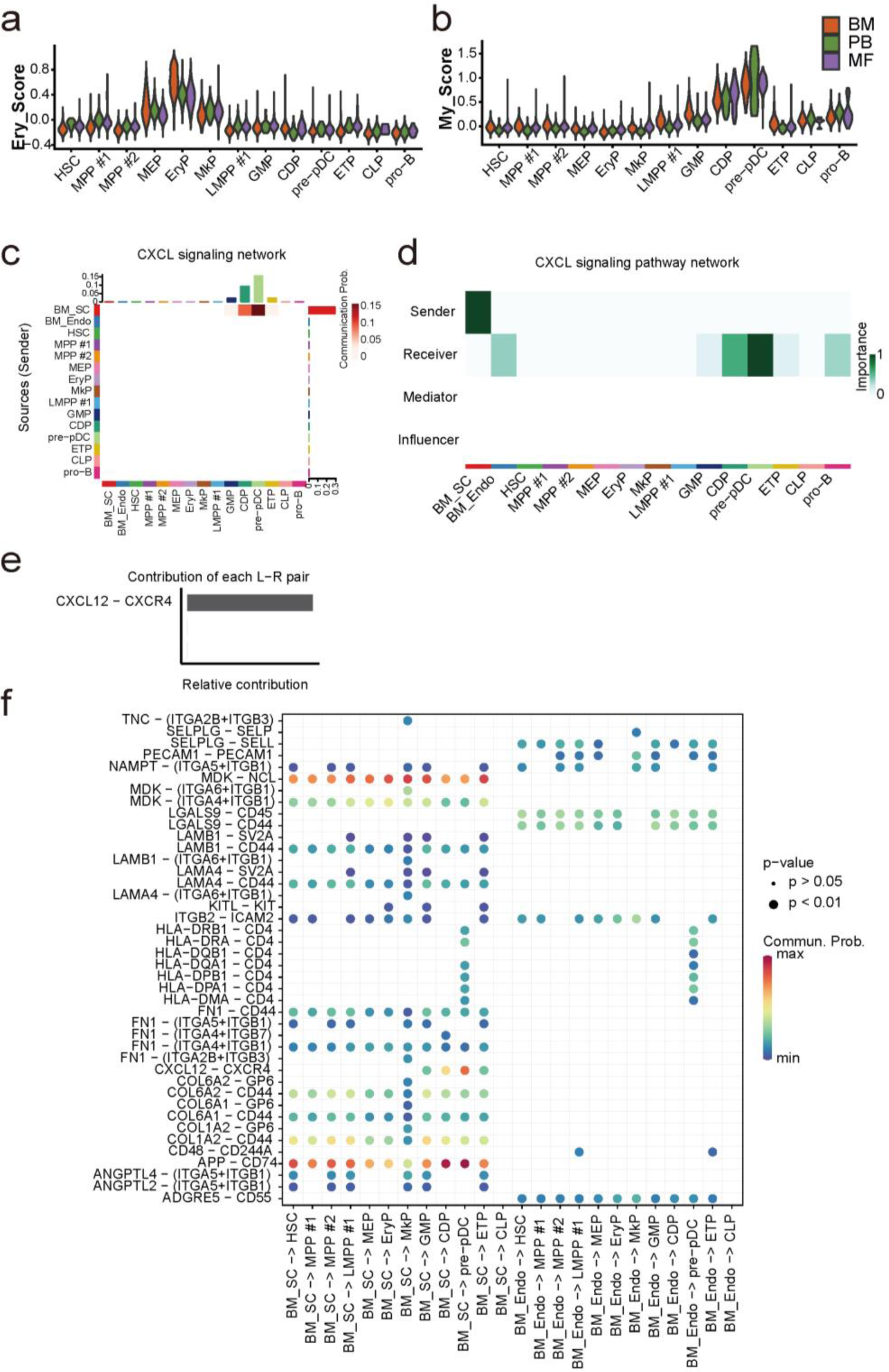
Cell signals from BM stromal subpopulations (BM_SC, BM_Endo) to HSPC subsets. **a-b**. Ery score **(a)** and My score **(b)** of HSPC subsets from BM, mPB, PB, and MF patients. **c-e**. cell-cell communication network (c), heatmap of ligand-receptor signaling (d) and contribution of ligand-receptor pairs (e) from BM_SC to HSPC subsets based on CXCL signaling. **f**. Interaction pairs from BM_SC, BM_Endo to HSPC subsets. Heatmap signaling between cell subsets in BM.

## Supplementary Tables

**Supplementary Table1.** The samples information of scRNA-seq data of human CD34+ cells.

**Supplementary Table2.** Primers of Cd99, B2m and Cd74.

